# Chromogranin A regulates gut permeability *via* the antagonistic actions of its proteolytic peptides

**DOI:** 10.1101/2020.09.19.304303

**Authors:** Elke M. Muntjewerff, Kechun Tang, Lisanne Lutter, Gustaf Christoffersson, Mara J.T. Nicolasen, Hong Gao, Gajanan D. Katkar, Soumita Das, Martin ter Beest, Wei Ying, Pradipta Ghosh, Sahar El Aidy, Bas Oldenburg, Geert van den Bogaart, Sushil K. Mahata

**Author notes:** **Correspondence should be addressed to**: Sushil K. Mahata, and Geert van den Bogaart,. shared co-first authorship (Elke M. Muntjewerff, Kechun Tang).

## Abstract

**Aim:** A ‘leaky’ gut barrier has been implicated in the initiation and progression of a multitude of diseases, e.g., inflammatory bowel disease, irritable bowel syndrome, celiac disease, and colorectal cancers. Here we show how pro-hormone Chromogranin A (CgA), produced by the enteroendocrine cells, and Catestatin (CST: hCgA_352-372_), the most abundant CgA-derived proteolytic peptide, affect the gut barrier.

**Methods:** Colon tissues from region-specific CST-knockout (CST-KO) mice, CgA-knockout (CgA-KO) and WT mice were analyzed by immunohistochemistry, ultrastructural and flowcytometry studies. FITC-dextran assays were used to measure intestinal barrier function. Mice were supplemented with CST or CgA fragment pancreastatin (PST: CgA_250-301_). The microbial composition of cecum was determined. CgA and CST levels were measured in blood of IBD patients.

**Results:** CST-KO mice displayed (i) elongated tight, adherens junctions and desmosomes similar to IBD patients, and (ii) gut inflammation. Consistently, plasma FITC-dextran measurements showed increased intestinal paracellular permeability in the CST-knockout mice. This correlated with a higher ratio of Firmicutes to Bacteroidetes, a dysbiotic pattern commonly encountered in various diseases. Supplementation of CST-knockout mice with recombinant CST restored paracellular permeability and reversed inflammation, whereas CgA-knockout mice supplementation with CST and/or PST in CgA-KO mice showed that intestinal paracellular permeability is regulated by the antagonistic roles of these two peptides: CST reduces and PST increases permeability.

**Conclusion:** The pro-hormone CgA regulates the intestinal paracellular permeability. CST is both necessary and sufficient to reduce permeability and primarily acts via antagonizing the effects of PST.

## Introduction

The intestinal tract harbors 10^3^-10^11^ bacteria/ml ^1^ that in aggregate can constitute >1 g of endotoxin ^2^. Endotoxins stimulate immune cells in the lamina propria, which can trigger mucosal inflammation ^3^. Such inflammation can result in luminal dysbiosis ^4^ and gut barrier defects ^5-8^. The gut barrier is comprised of a surface mucus layer which separates intact bacteria and large particulates from the mucosa ^9^ and the colonic epithelium which provides a physical and immunological barrier separating the body from the luminal contents (e.g., dietary antigens, a diverse intestinal microbiome, and pathogens) ^10-12^. Mucosal homeostasis is maintained by the delicate and complex interactions between epithelial cells, immune cells, and luminal microbiota ^10^.

The gastrointestinal epithelium not only forms the body’s largest interface with the external environment, but also establishes a selectively permeable epithelial barrier; it allows nutrient absorption and waste secretion while preventing the entry of luminal contents. The epithelial barrier is regulated by a series of intercellular junctions between polarized cells: an apical tight junction (TJ), which guards paracellular permeability, the subjacent adherens junction (AJ) and desmosomes; the AJs and desmosomes do not seal the paracellular space but provide essential adhesive and mechanical properties that contribute to paracellular barrier functions ^11^.

The permeability of the barrier is regulated by many factors such as the immune system. For example, tumor necrosis factor (TNF)-α increases the flux of larger molecules, such as proteins and other macromolecules, through the leak pathway, and interleukin (IL)-13 selectively enhances the flux of small molecules and ions via the pore pathway ^5^. TNF-α plays crucial roles in the Caveolin-1-mediated internalization of Occludin (OCLN), which elevates gut permeability. Conversely, overexpression of OCLN alleviates the cytokine-induced increase in gut permeability ^6^. Interferon (IFN)-*γ* increases gut permeability by reducing the expression of Tight junction protein ZO-1 (TJP1) and OCLN as well as by modulating actin-myosin cytoskeleton interactions with TJ proteins ^7^. The simultaneous presence of TNF-α and IFN-*γ* deteriorates intestinal integrity by dissociating TJ proteins ^8^.

Here we show that the permeability of the gut barrier is regulated by the pro-hormone Chromogranin A (CgA). CgA is an acidic secretory proprotein ^13^ that is abundantly expressed in enteroendocrine cells (EECs) of the gut and serves as a common marker for the EECs of the gut ^14^. Originally, CgA was identified as a cellular packaging factor ^15-17^. However, the pro-hormone CgA is proteolytically cleaved, both intracellularly as well as extracellularly after its secretion, giving rise to seven bioactive peptides that exert a wide variety of regulatory functions among the metabolic ^18-21^, cardiovascular ^22-25^ and immune systems ^26-30^. Among these bioactive peptide hormones are catestatin (CST: hCgA_352-372_) ^31,32^ and pancreastatin (PST: hCgA_250-301_) ^33^. Blood plasma levels of both intact CgA ^34-36^ and CST ^37^ are increased in patients with inflammatory bowel disease (IBD). We show that CST is abundantly generated in the gut. Unexpectedly, we found that CgA regulates the paracellular permeability of the gut via the antagonistic actions of its two peptides, PST and CST; while PST is a pro-inflammatory, pro-obesigenic and anti-insulin peptide ^20^ which loosens the barrier, CST acts as anti-inflammatory, antiobesigenic, anti-microbial, and proinsulin peptide ^25,27,29^ that appears to be necessary and sufficient to neutralize the actions of PST and tighten the barrier. Using knockout (KO) mice in conjunction with recombinant proteins, we show that CST exerts its actions *via* counteracting PST.

## Results

### CgA is extensively processed to CST

To study the role of CgA and its peptides in the regulation of the gut barrier, we used systemic CgA-KO (deletion of exon 1 and ∼1.5 kbp proximal promoter, thereby completely inactivating the *Chga* allele) ^22^ and CST-KO mice (the 63 bp CST domain of CgA was deleted from Exon VII of the *Chga* gene) ^38^. CST-KO mice show low grade inflammation in the heart, pancreas, and liver and have an elevated blood pressure, are obese and suffer from insulin resistance ^27,38^. We analyzed the function and morphology of the gut barrier in CST-KO mice. We used transmission electron microscopy (TEM) to resolve the ultrastructure of EECs and to evaluate the subcellular compartment where hormones are stored and released, i.e., dense core vesicles (DCVs). While DCV numbers were reduced in both CgA-KO and CST-KO mice, DCV areas were smaller in CgA-KO mice but larger in CST-KO mice (Fig. 1A&B). WT colon showed a lower concentration of CgA (average 0.019 nmol/mg protein) than of CST (average 0.37 nmol/mg protein), indicating extensive processing of CgA to CST (Fig. 1C). As anticipated, in CST-KO mice the levels of CgA in the colon were similar to that in WT mice, but those of CST were undetectable (Fig. 1C). Consistent with enhanced processing of CgA, we found increased expression of the proteolytic enzymes Cathepsin L (*Ctsl*, which cleaves CST from CgA) and Plasminogen (*Plg*, the precursor of plasmin, which is involved in CgA processing) in CST-KO and CgA-KO, respectively (Fig. 1D) ^39,40^.

**Fig. 1.**
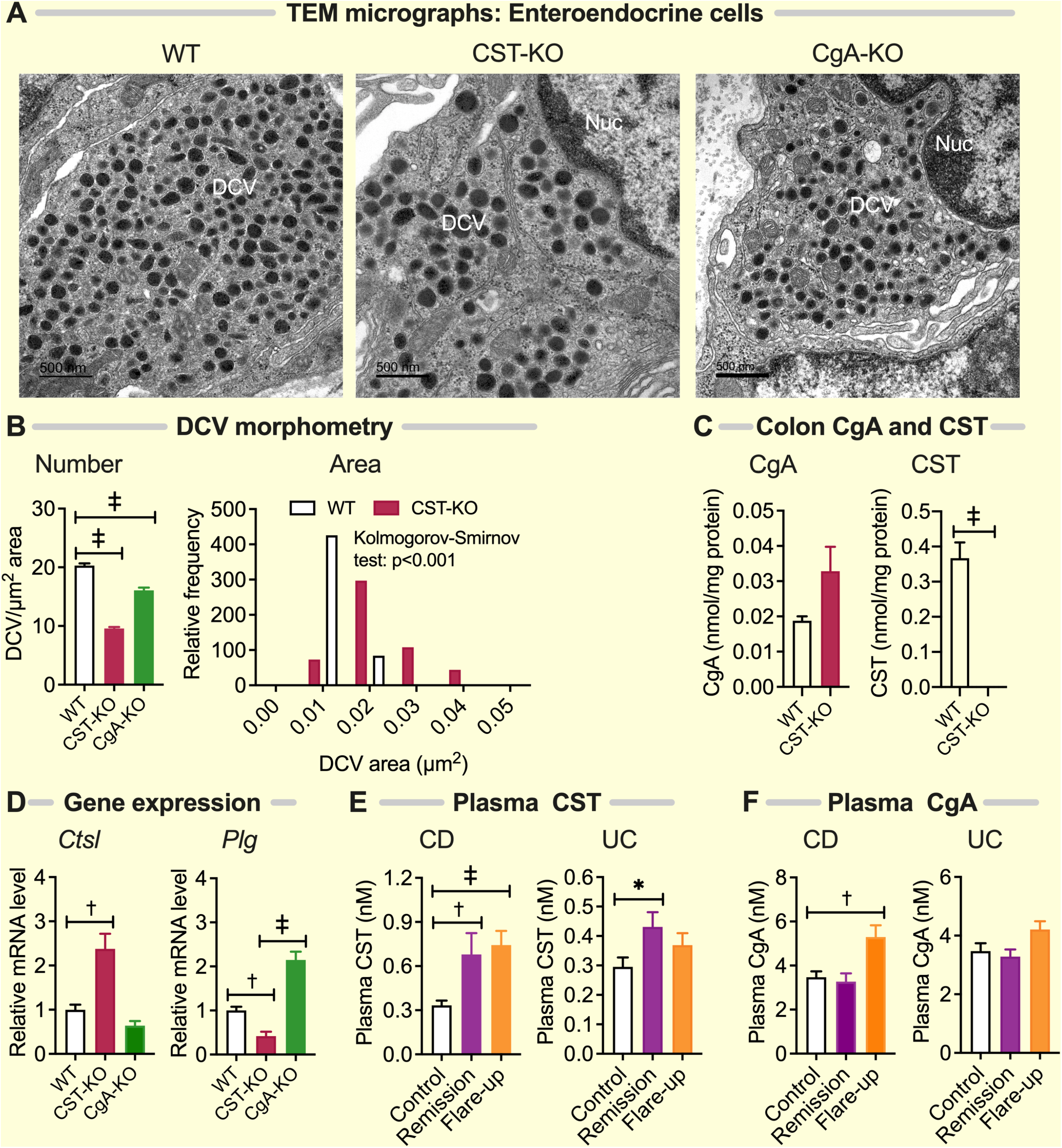
CgA is efficiently processed to CST in the colon and CST levels correlate with Crohn’s disease activity. **(A)** TEM micrographs (n=3) showing enteroendocrine cell DCV in colon of WT, CgA-KO and CST-KO mice, **(B)** fewer DCV in CgA-KO and CST-KO mice (1-way ANOVA) but enlarged DCV in CST-KO mice and smaller DCV in CgA-KO mice (>500 vesicles from 30 micrographs) (Kolmogorov-Smirnov test). DCV, dense core vesicles; Nuc, nucleus. **(C)** CgA protein and CST peptide levels in colon of WT and CST-KO mice. CgA levels were comparable between WT and CST-KO mice. CST level was markedly higher in WT mice and was undetectable in CST-KO mice (n=6; Welch’s *t* test). **(D)** Expression of proteolytic genes (n=7). *Ctsl* and *Plg* are differentially expressed in colon of CgA-KO and CST-KO mice (n=8; 1-way ANOVA). **(E)** CgA and CST levels in EDTA-plasma of healthy donors (Control) and Crohn’s disease patients (N=89) in remission or flare-up (2-way ANOVA). Ns, not significant; †, p<0.01; ‡, p<0.001.

Because the processing of CgA to CST is compromised in several diseases ^41-43^, we determined plasma CgA and CST levels in patients with Crohn’s disease and with ulcerative colitis, both with active disease and in remission (Fig. 1E&F). In line with previous findings ^34-36^, we found higher CgA levels in plasma of patients with active Crohn’s disease but not in patients in remission. In contrast, we found higher plasma CST levels in Crohn’s disease regardless of disease activity, indicating that the processing of CgA to CST is more efficient in active disease. Patient characteristics are provided in Supplemental Table S1&S2.

### CST-KO mice have a ‘leaky’ gut

Ultrastructural evaluation of the gut epithelium revealed that the TJs, AJs and desmosomes are significantly different in CST-KO mice compared to wild-type (WT): the TJs, AJs and desmosomes were elongated, as also observed in IBD patients (Fig. 2A, C-E, Fig. 3H-L). We determined whether this altered morphology was caused by an altered expression of proteins involved in these epithelial junctions. Gene expression studies showed a trend towards mRNA expression of multiple TJ-markers such as Occludin (*Ocln*), MARVEL domain-containing protein 2 (*Marveld2*, also called Tricellulin), Junctional adhesion molecule A (JAM-A; *F11r*) Tight junction protein ZO-1 (*Tjp1*) ^44^ and the desmosomal protein Desmoglein 2 (*Dsg2*), which is required for maintenance of intestinal barrier function ^45^. We observed concomitant elevated mRNA expression of the genes coding for TJ components Claudin 1 and 2 (*Cldn1* and *Cldn2*), TJ protein ZO-2 and 3 (*Tjp2* and *Tjp3*), and AJ genes coding for alpha-E-Catenin (*Ctnna1*), Cadherin-associated protein (*Ctnnb1*), and Cadherin 1 (*Cdh1*) (Fig. 2F&G). Primer sequences are provided in Supplemental Table S3. For Occludin, ZO-1, alpha-E-Catenin, Cadherin 1, Claudin 1, we did not observe significantly altered protein levels. However, we did find expression at both the mRNA and protein levels of *Cldn2* (Fig. 2F, Fig. 3A&G), a specific claudin that is known to increase the paracellular flux of cations and water through the pore pathway ^46^.

**Fig. 2.**
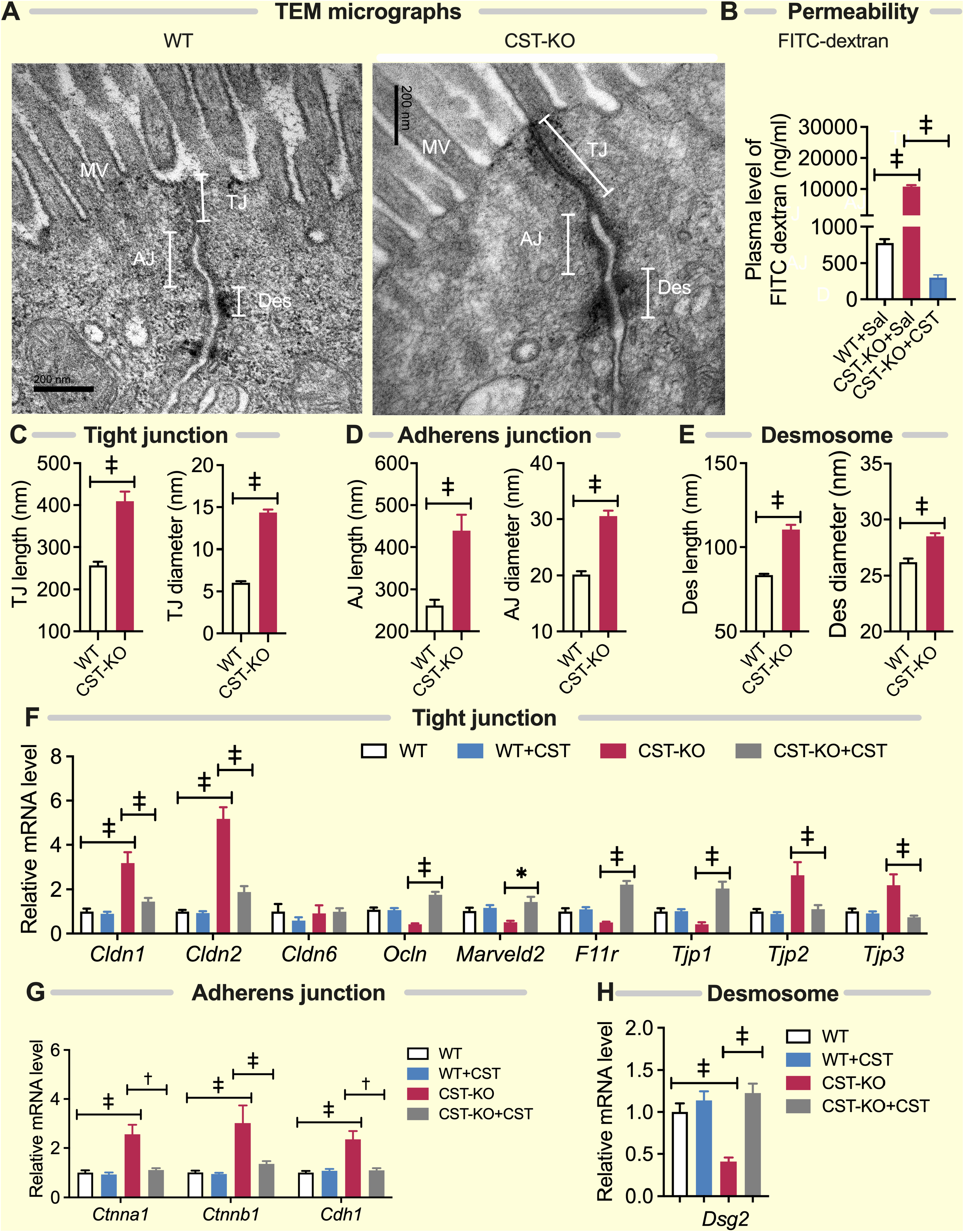
Impaired epithelial paracellular barrier function in CST-KO mice. **(A)** TEM micrographs (n=3) showing tight junction (TJ), adherens junction (AJ), and desmosomes (Des). MV, microvillus. **(B)** Increased gut permeability as measured by FITC-dextran level in CST-KO compared to WT mice (n=8; Welch’s *t* test). (**C-E**) Morphometrical analyses (n=20-54) showing increased length and diameter of TJ **(C)**, AJ **(D)** and Des **(E)** in CST-KO mice (Welch’s *t* test). **(F-H)** Differential expression of genes in TJ (*Cldn1, Cldn2, Cldn6, Ocln, Marveld2, F11r, Tjp1, Tjp2, and Tjp3*), AJ (*Ctnna1, Ctnnb1*, and *Cdh1*) and Des (*Dsg2*) in colon of CST-KO mice compared to WT mice. Supplementation of CST-KO mice with CST reversed the expression of the above genes (n=8; 2-way ANOVA or Welch’s *t* test). *, p<0.05; †, p<0.01; ‡, p<0.001.

**Fig. 3.**
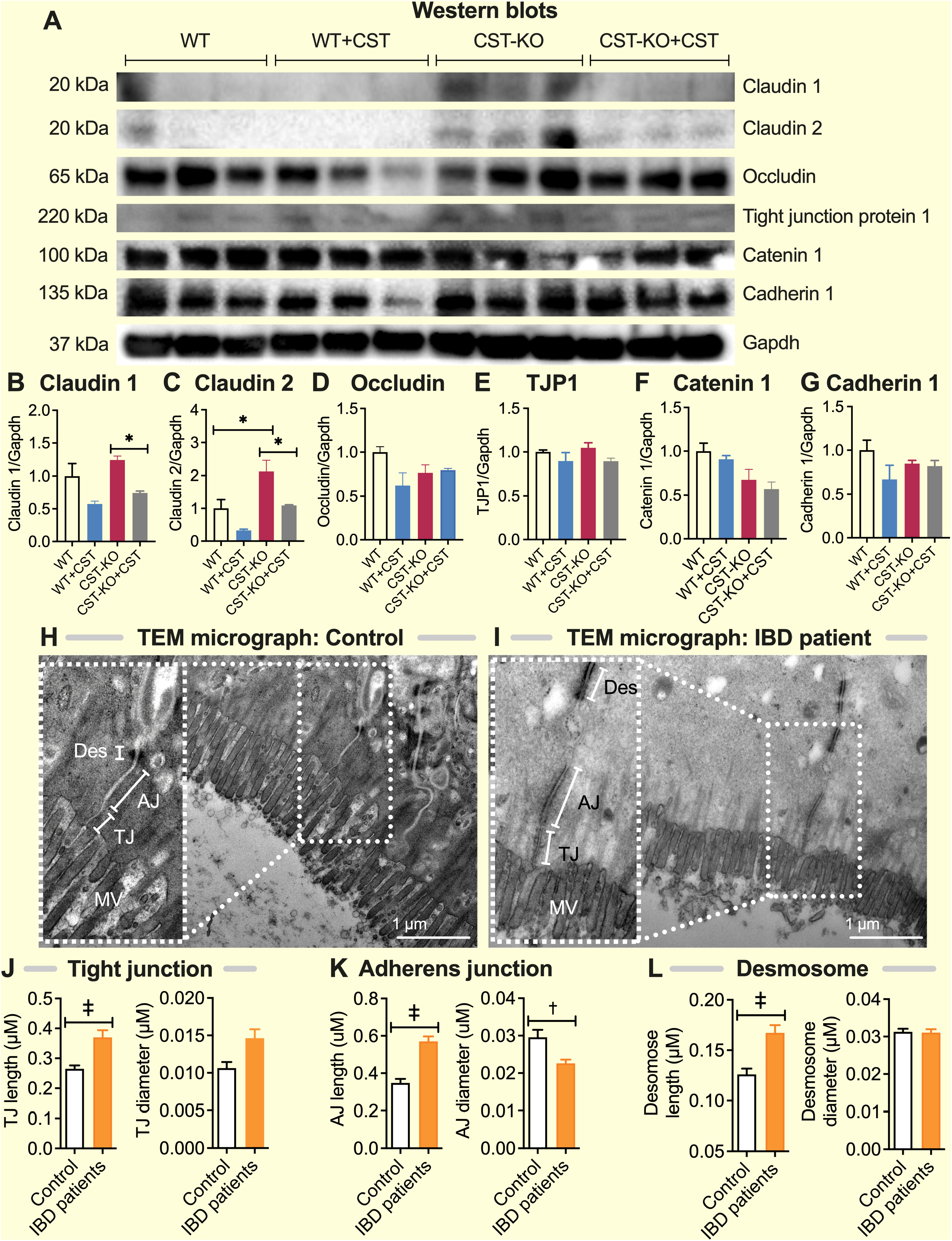
Elevated levels of Claudin-2 in colon of CST-KO mice and elongated epithelial junctions in IBD patients. **(A)** Western blots and **(B)** densitometric analyses of Claudin-1 **(B)**, Claudin-2 **(C)**, Occludin **(D)**, Tjp1 **(E)**, Catenin 1 **(F)**, and Cadherin 1 (**G**). (**H&I**) TEM micrographs (n=3) showing tight junction (TJ), adherens junction (AJ), and desmosomes (Des) in healthy control and IBD patients. Microvillus (MV). (**J-L**) Morphometrical analyses (n=20-54) showing increased lengths and diameter of TJ (**J**), AJ (**K**) and Des (**L**) in IBD patients. *, p<0.05; †, p<0.01; ‡, p<0.001.

In order to assess the functional consequence of the altered cellular junctions, we determined the flux through the leak pathway, which is responsible for the paracellular movement of larges molecules, including microbial proteins and other macromolecules. Compared to WT mice, we observed a more than 10-fold increase in leakage of plasma FITC-dextran (4 kDa) in the CST-KO mice (Fig. 2B). This confirms that the intestine in CST-KO mice has a strongly diminished paracellular barrier via the leak pathway, although the large magnitude of leakage might even suggest direct epithelial damage.

### CST-KO mice and IBD patients show comparable ultrastructural changes in tight junctions, adherens junctions, and desmosomes

Ultrastructural analyses of colonic mucosa revealed increased lengths of TJ, AJ, and desmosomes in IBD patients compared to healthy controls (Fig. 3H-L). Unlike CST-KO mice, IBD patients displayed a decrease in AJ diameter (Fig. 3H&K).

### The leaky gut in CST-KO mice is accompanied by mucosal inflammation and dysbiosis

Because inflammation regulates the epithelial cellular junctions ^5-8^ and an increased gut permeability is often associated with mucosal inflammation, we investigated the immune cell populations in the colons of CST-KO mice by histochemical analysis (Fig. 4A-C). Indeed, the colon of CST-KO mice displayed signs of mucosal inflammation, such as elevated immune cell infiltration (Fig. 4D&E), increased fibrosis (Fig. 5D&E), and increased apoptosis (Fig. 6A). In addition, there was increased abundance of macrophages (CD45^+^11b^+^Emr1^+^) and helper CD4^+^ T cells in the colon of CST-KO mice as confirmed by flow cytometry (Fig. 4F). Moreover, CST-KO mice showed increased expression of the following proinflammatory genes: TNF-α (*Tnfa*), IFN-γ (*Ifng*), Chemokine CXC motif ligand 1 (*Cxcl1* aka *Kc/Gro1*), Chemokine CC motif ligand 2 (*Ccl2* aka *Mcp1*), Egg-like module-containing mucin-like hormone receptor 1 (*Emr1* aka *F4/80*), Integrin alpha-M (*Itgam* aka *Cd11b*), and Integrin alpha-X (*Itgax* aka *Cd11c*) (Fig. 4G). Increased expression of *Tnfa, Ifng, Cxcl1*, and *Ccl2* genes correlated with increased protein levels (Fig. 5A). Upregulation of proinflammatory cytokines is in line with prior work demonstrating a positive correlation of IFN-γ ^47^ and TNF-α ^44^ with intestinal leakiness. The decreased expression of anti-inflammatory cytokines, such as interleukin 4 (*Il4*) and interleukin 10 (*Il10*), in CST-KO mice is also in congruence with the reported role of anti-inflammatory cytokines in gut permeability (Fig. 5C) ^48^. The decreased *Il10* mRNA levels correlated with a decreased protein level of IL-10 (Fig. 5C). Furthermore, the expression of *S100a9* gene (forming Calprotectin in complex with S100a8) was increased in colon of CST-KO mice and Calprotectin protein levels were increased in plasma, colon and feces of CST-KO mice (Fig. 5B); increased Calprotectin, a protein secreted by activated neutrophils and macrophages, has been shown to be associated with increased intestinal permeability ^49^. The increased inflammation in the gut of CST-KO mice is consistent with the known anti-inflammatory roles of CST ^29,50^ and with our previous observation of low-grade systemic inflammation in the CST-KO mice ^27^.

**Fig. 4.**
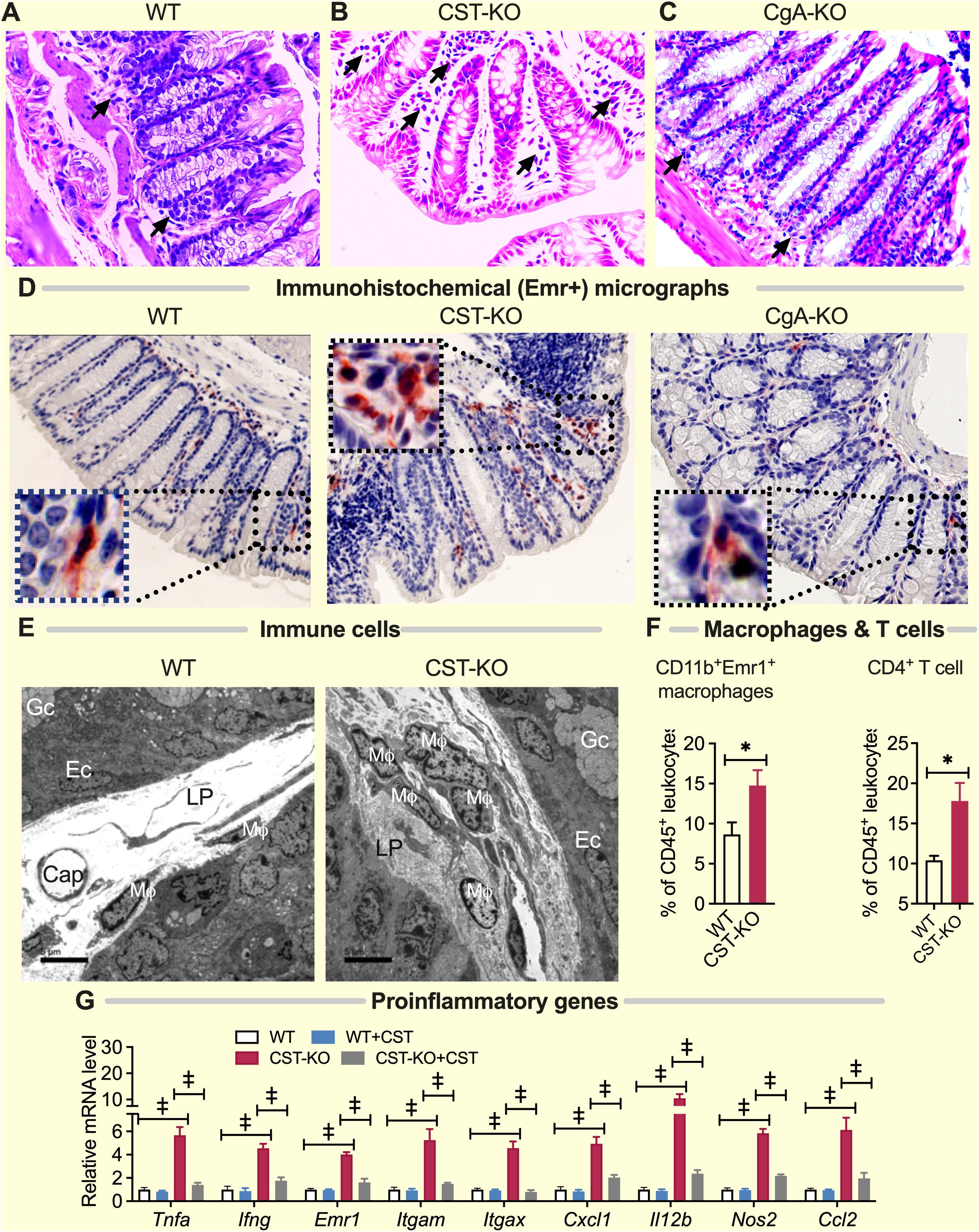
Increased infiltration of immune cells in colon of CST-KO mice. **(A-C)** hematoxylin and eosin-stained histological micrographs of colon in WT **(A)**, CST-KO **(B)**, and CgA-KO **(C)** mice. Arrows indicate immune cells. **(D)** Immunohistological micrographs showing increased infiltration of Emr^+^ cells in CST-KO mice. **(E)** TEM micrographs (n=3) showing increased number of immune cells in colon of CST-KO mice. Mϕ, macrophage; Cap, capillary; Ec, enterocytes; Gc, goblet cells; LP, lamina propria. **(F)** Flow cytometry study (n=5-10) shows increased number of macrophages (CD45^+^CD11b^+^Emr1^+^) and T cells (CD45^+^CD4^+^) in colon of CST-KO mice (n=5-10; Welch’s *t* test). **(G)** Increased expression of proinflammatory genes (*Tnfa, Ifng, Cxcl1, Ccl2, Emr1, Il12b, Nos2, Itgam*, and *Itgax*), which returned to WT levels after treatment with CST (n=8; 2-way ANOVA). (n=8; 2-way ANOVA). *, p<0.05; †, p<0.01; ‡, p<0.001.

**Fig. 5.**
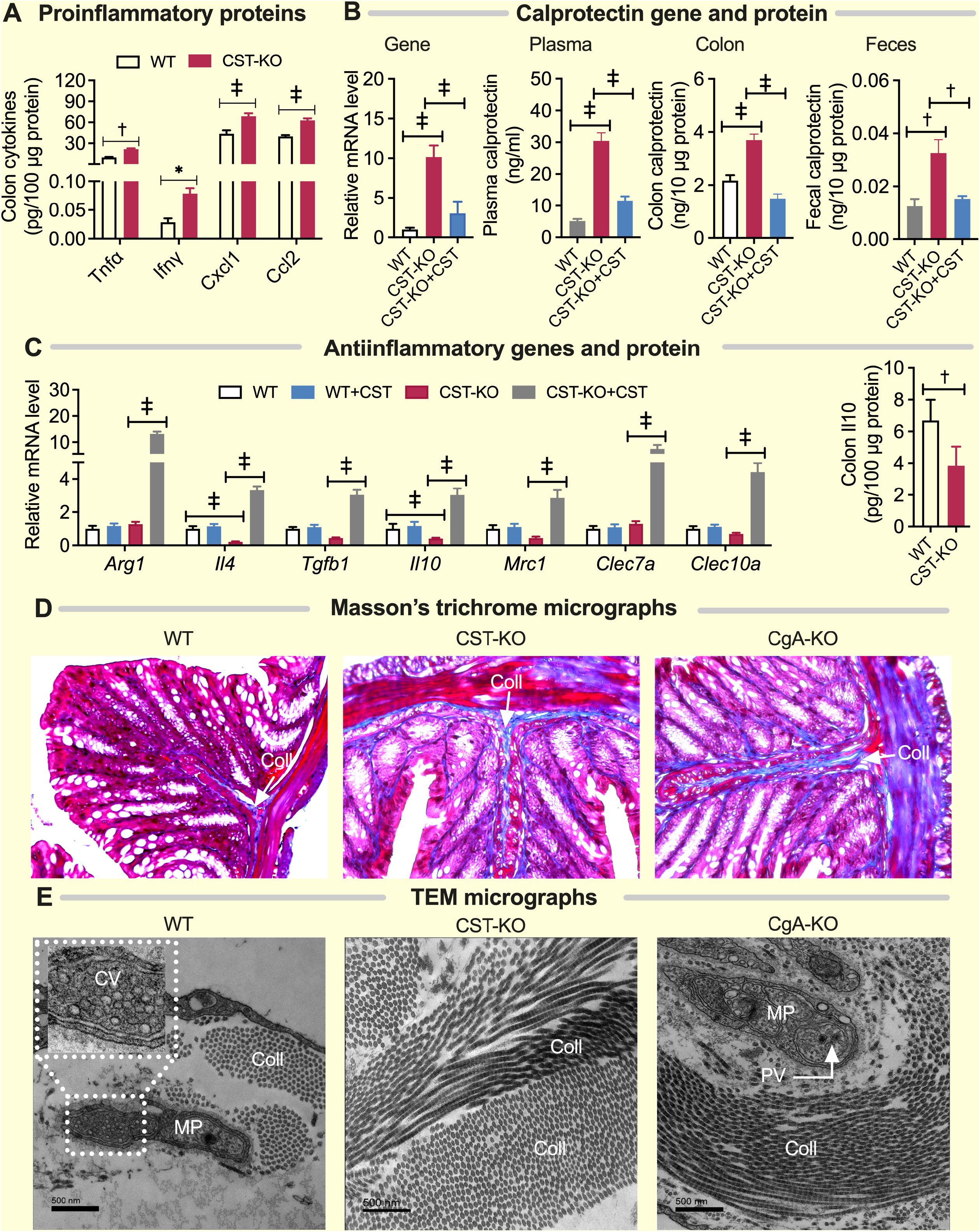
Elevated levels of pro-inflammatory proteins and fibrosis in colon of CST-KO mice. **(A)** Increased expression of proinflammatory proteins (Tnfα, Ifnγ, Cxcl1, and Ccl2) in colon of CST-KO mice (n=8; 2-way ANOVA). **(B)** Increased expression of proinflammatory Calprotectin gene *s100A9* in colon and Calprotectin protein in plasma, colon, and feces of CST-KO mice, which were reversed upon administration of CST for 15 days (n=5; 1-way ANOVA). **(C)** Decreased expression of anti-inflammatory genes (*Arg1, Il4, Tgfb1, Il10, Mrc1, Clec7a*, and *Clec10a*) (n=8; 2-way ANOVA), and anti-inflammatory protein (Il10) (n=8; Welch’s *t* test) in colon of CST-KO mice. CST treatment increased expression of anti-inflammatory genes in CST-KO mice. **(D)** Masson’s trichrome-stained histological micrographs showing increased fibrosis in CST-KO and CgA-KO colon. **(E)** TEM micrographs showing increased presence of collagen fibers in CST-KO and CgA-KO mice. Coll, collagen; CV, clear cholinergic vesicles; MP, Meissner’s plexus; PV, dense core peptidergic vesicles. *p<0.05; **p<0.01; ***p<0.001.

**Fig. 6.**
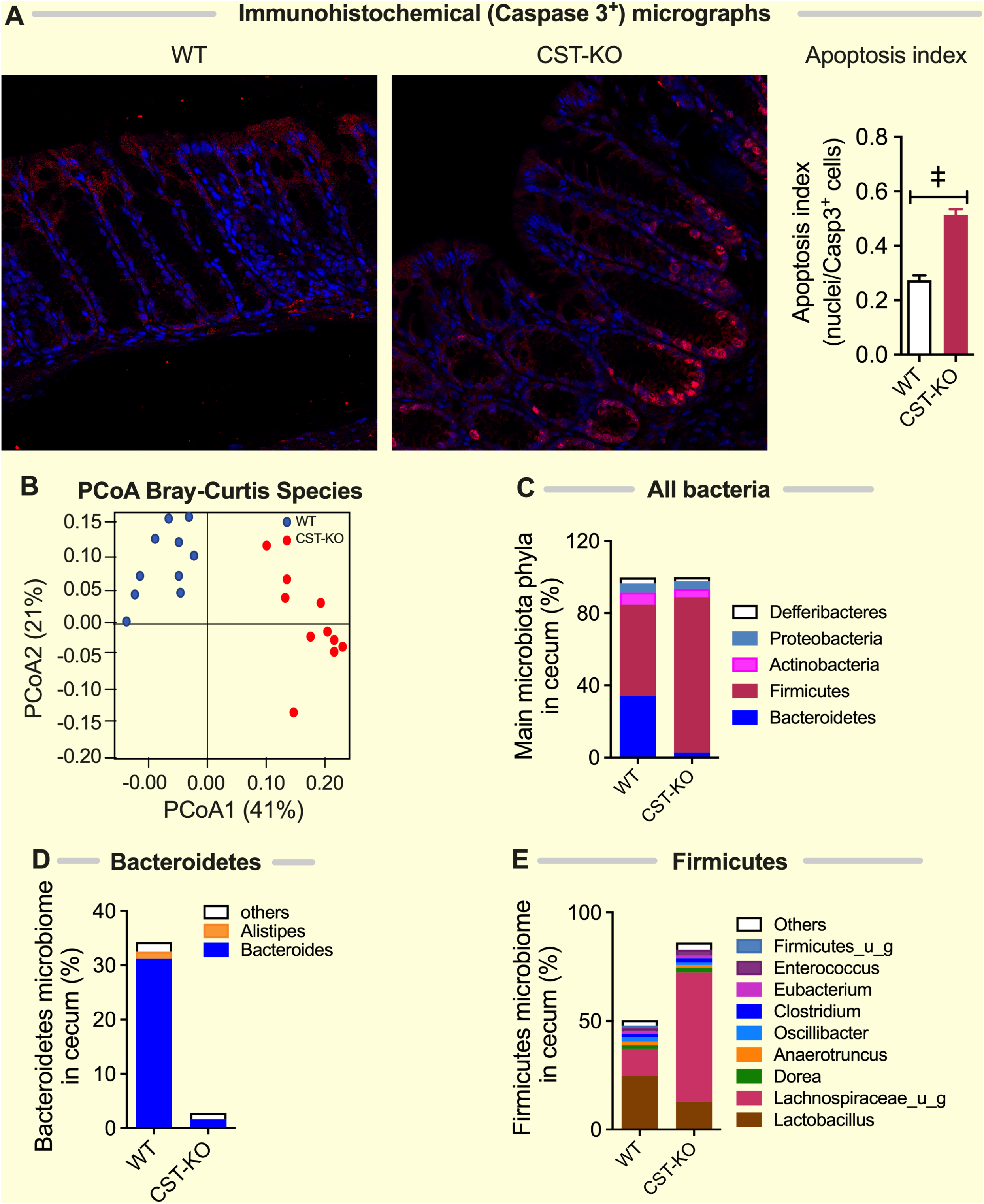
Increased apoptosis and ratio of Firmicutes to Bacteriodetes in gut of CST-KO mice. **(A)** increased numbers of Caspase-3 positive apoptotic cells in colon of CST-KO mice. **(B)** Principal CoordiNATE Analysis (PCoA) of microbiome data using Bray-Curtis distance of species composition and abundance in individual WT (blue) and CST-KO (red) mice. **(C-E)** Microbiota composition of cecum of WT and CST-KO mice for main bacteria phyla **(C)** and the geni within the *Bacteriodetes* **(D)** and *Firmicutes* **(E)**.

Because a leaky gut may either cause or be the consequence of altered luminal microbes ^51^, we investigated whether the microbial population in CST-KO mice was altered. We found that the microbiome in CST-KO mice was indeed quite different in composition than its WT littermates (Fig. 6B-E&7). Most prominently, CST-KO mice showed a higher ratio of Firmicutes to Bacteroidetes (Fig. 6C-E), which is opposite to the decreased ratio observed in WT mice intra-rectally infused with CST ^52^, and supports the idea that a low ratio of Firmicutes to Bacteriodetes could be beneficial to dampen intestinal inflammation ^52^.

That the leaky gut of CST-KO mice is inflamed and harbors dysbiotic luminal contents implies that CST is required for maintaining gut barrier integrity and preventing dysbiosis and gut inflammation. This is important because a “leaky gut” ^53^, dysbiosis ^54^, and hyperactivated immunity ^55^ are all well known to play a role in IBD. It is also consistent with others’ observation that administration of exogenous CST can ameliorate colitis in mice ^55^.

### CST is sufficient to tighten the leaky gut barrier in CST-KO mice and reduce inflammation

To test whether CST is sufficient to regulate gut permeability, we supplemented CST-KO mice with recombinant CST (intraperitoneal injections of 2 µg/g body weight once daily for at least 15 days), which reversed paracellular leakiness of FITC-dextran and the associated abnormalities such as: (i) reduced plasma FITC-dextran even lower than WT level (Fig. 2B), (ii) increased mRNA expression of *Ocln, Marveld2, F11r, Tjp1, and Dsg2* (Fig. 2F&H), (iii) decreased mRNA expression *of Cldn1, Cldn2, Tjp2, Tjp3, Ctnna1, Ctnnb1*, and *Cdh1*, as well as increased protein expression of Claudin-2 (Fig. 2F&G, 3A&C), (iv) decreased mRNA expression of proinflammatory genes (*Tnfa, Ifng, Cxcl1, Ccl2, Emr1, Il12b, Nos2, Itgam*, and *Itgax)* (Fig. 4G), (v) increased expression of anti-inflammatory genes (*Arg1, Il4, Tgfb1, Il10, Mrc1, Clec7a*, and *Clec10a*) (Fig. 5C), (vi) decreased expression of Calprotectin gene (subunit *S100a9)* and protein levels (Fig. 5B). These findings demonstrate that CST reduces intestinal inflammation, and that CST is not only required for the integrity of the paracellular gut barrier, but is also sufficient to maintain the integrity of this barrier during homeostasis.

### The paracellular permeability of the gut is intact in CgA-KO mice

Next, we carried out the gut barrier studies in CgA-KO mice, which not only lack CST, but also lack all other peptides derived from CgA. Ultrastructural studies showed no structural changes in TJ, AJ and desmosomes in CgA-KO mice compared to WT mice (Fig. 8A). To our surprise, we found that despite the absence of CST in these mice, the gut permeability for FITC-dextran was not increased (Fig. 8B); if anything, the paracellular barrier was even tighter than WT control mice. This decreased permeability was not accompanied by changes in CgA-KO mice in genes coding for components of TJ, AJ and desmosomes, except that *Ocln* and *Dsg2* gene expression were decreased in CgA-KO mice compared to WT mice (Fig. 8C). Likewise, proinflammatory gene and protein expressions were not affected in CgA-KO mice (Fig. 8D-E). In line with this, histology and EM revealed some fibrosis, but no excessive immune cell infiltration in the colon of CgA-KO mice (Fig. 5D&E). In contrast to the CST-KO mice, the CgA-KO mice even showed increased colonic expression of anti-inflammatory genes and proteins (Fig. 8F-G). The decreased colon levels of Calprotectin were also consistent with decreased gut permeability (Fig. 8H). Compared to CST-KO mice, Firmicutes and Bacteroidetes bacteria populations in CgA-KO mice were more comparable to WT mice (Fig. 7&9A).

**Fig. 7.**
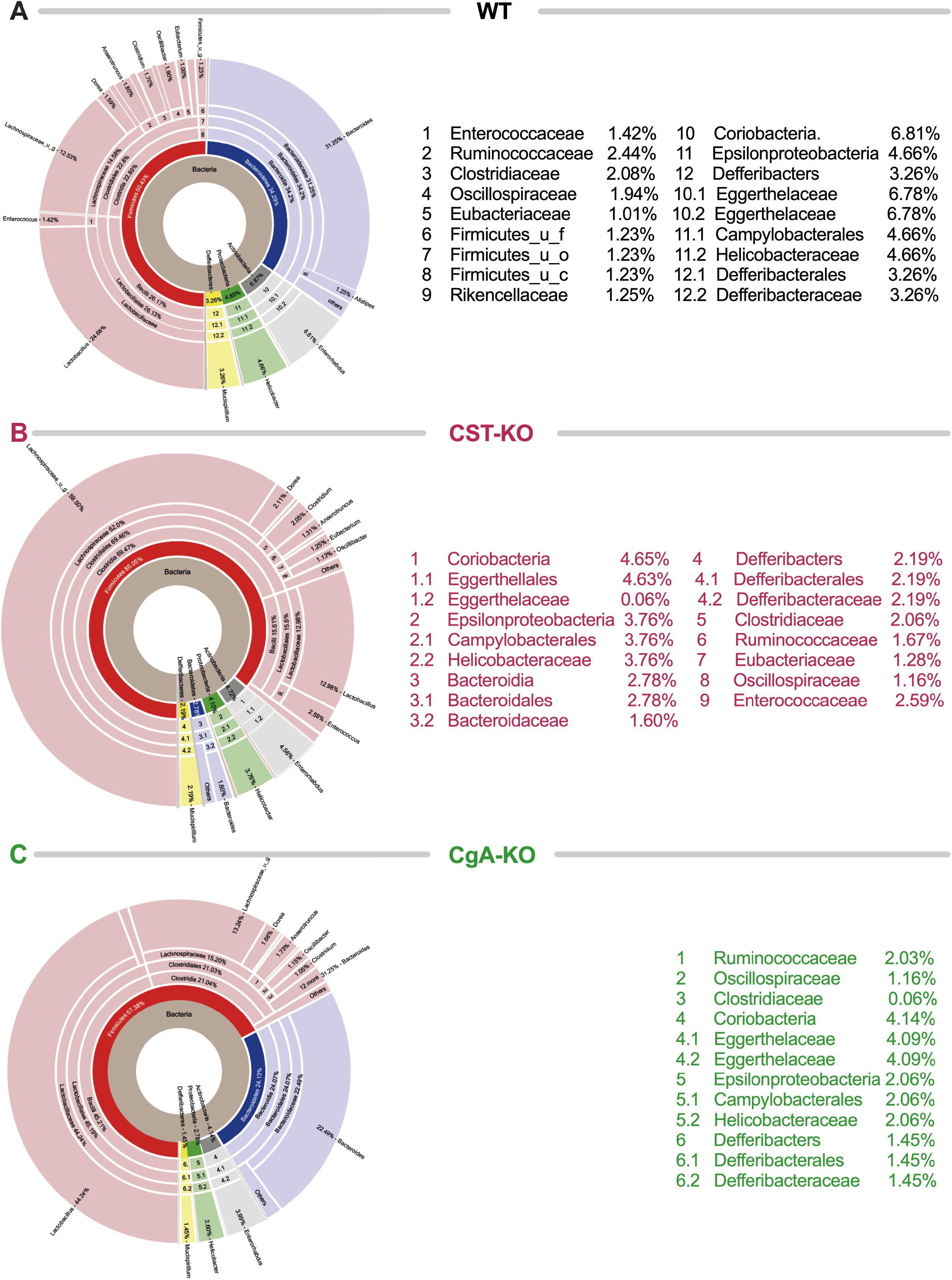
Detailed microbial composition of WT, CST-KO, and CgA-KO mice. Graphs show complete microbiota composition in cecum of WT **(A)**, CST-KO **(B)**, and CgA-KO **(C)** mice. Circles from inside to outside show kingdom, phylum, class, order, family and genus.

**Fig. 8.**
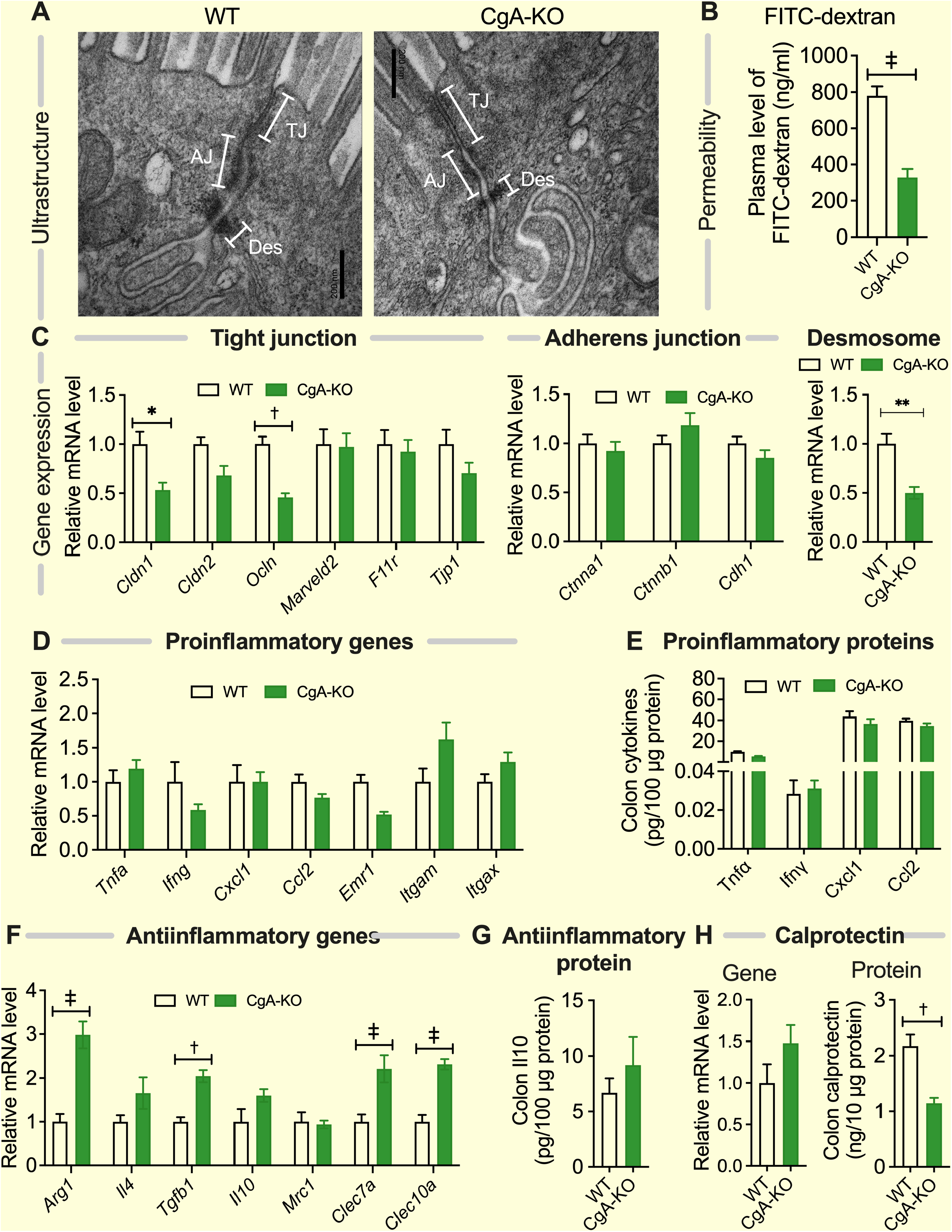
Increased epithelial paracellular barrier function and lower inflammation in CgA-KO mice. **(A)** Electron microscopy micrographs of colon of WT and CgA-KO mice. TJ: tight junction. AJ: adherens junction. Des: desmosomes. **(B)** Gut permeability as measured by FITC-Dextran plasma level in WT and CgA-KO mice (n=7; Welch’s *t* test). **(C)** Relative mRNA expression of genes in tight junction (*Cldn2, Ocln, Marveld2, F11r*, and *Tjp1*) (n=8, 2-way ANOVA), adherens junction (*Ctnna1, Ctnnb1*, and *Cdh1*) (n=8; 2-way ANOVA), and desmosome (*Dsg2*) (n=8; Welch’s *t* test) in colon of WT and CgA-KO mice. Comparable expression of **(D)** proinflammatory genes (*Tnfa, Ifng, Cxcl1, Ccl2, Emr1, Itgam*, and *Itgax*), and **(E)** proinflammatory proteins (Tnfα, Ifnγ, Cxcl1, and Ccl2) in colon of WT and CgA-KO mice (n=8, 2-way ANOVA). Increased expression of **(F)** anti-inflammatory genes (*Arg1, Tgfb1, Clec7a, and Clec10a*) and comparable expression of anti-inflammatory genes (*Il4, Il10*, and *Mrc1*), and **(G)** anti-inflammatory protein (Il10) in colon of WT and CgA-KO mice (n=8; 2-way ANOVA or Welch’s *t* test). **(H)** Comparable expression of *s100a9* gene and decreased expression of calprotectin protein in CgA-KO colon (n=5; Welch’s *t* test). †, P<0.01; ‡, P<0.001.

**Fig. 9.**
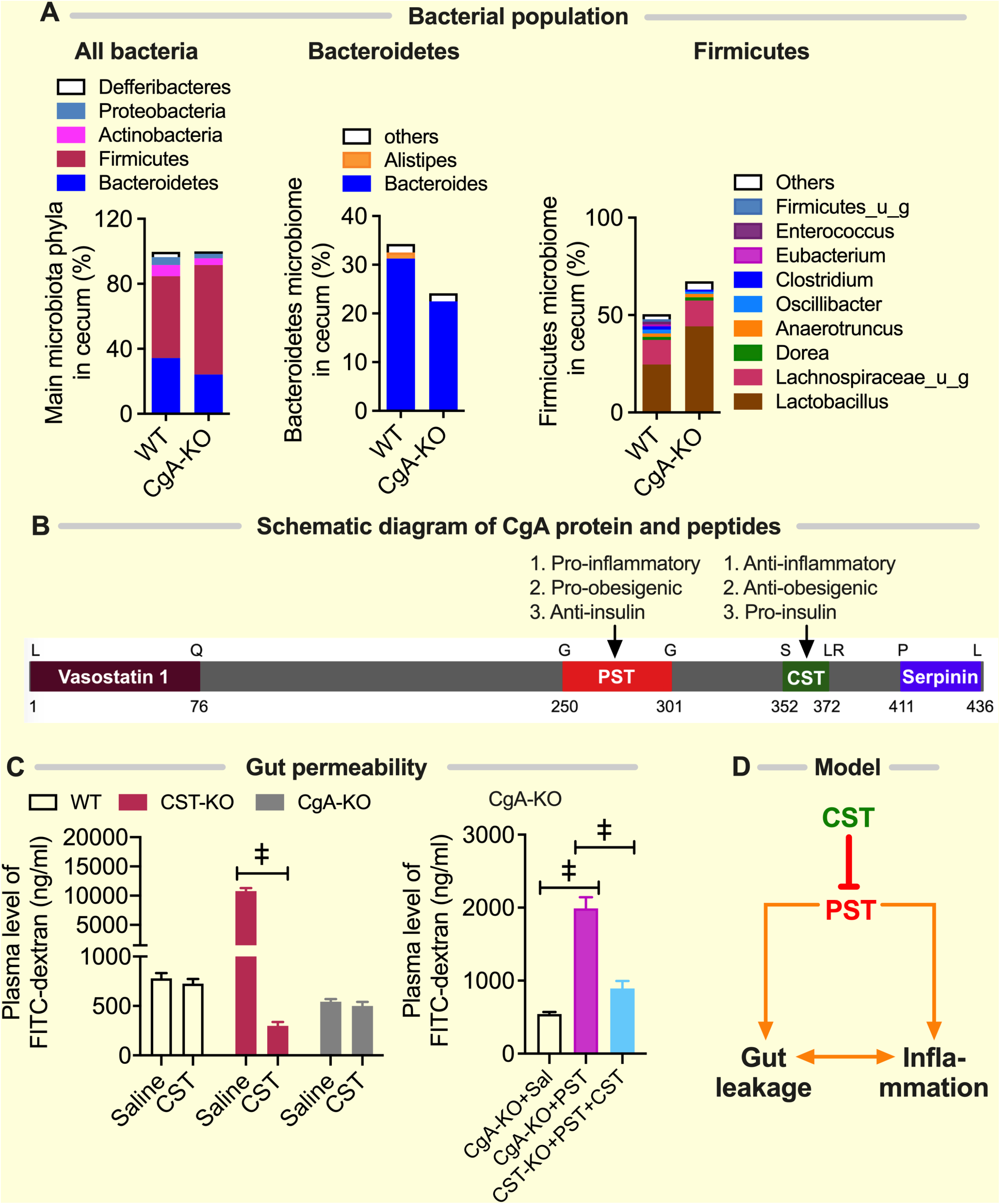
Altered microbiome composition in CgA-KO mice and CST and PST oppositely regulate gut permeability. **(A)** Microbiota composition of cecum of WT and CgA-KO mice for main bacteria phyla and the geni within the Bacteriodetes and Firmicutes. **(B)** Scheme of CgA indicating the positions of the peptide products vasostatin 1, PST, CST and serpinin. **(C)** Gut permeability as measured by FITC-Dextran plasma levels in WT, CST-KO and CgA-KO mice treated with saline or CST (n=8; 2-way ANOVA) and in CgA-KO mice treated with PST or a combination of PST and CST (n=8; 1-way ANOVA). **(D)** Model of gut barrier regulation by CST and PST. ‡, P<0.001.

These findings suggest that CST is not essential in the absence of CgA. Because a leaky and inflamed gut was seen in the CST-KO mice which specifically lack the proteolytic fragment of CST, but not in CgA-KO mice which lack CST and all other peptides that are generated proteolytically from CgA, these findings raised the possibility that CST may be required to antagonize the actions of some other peptide hormone that is also a product of CgA.

### Gut permeability is regulated by the antagonistic effects of two CgA peptides: CST and Pancreastatin (PST)

Among the various proteolytic peptides, we hypothesized that CST could be acting to balance the impact of PST, another peptide fragment of CgA (Fig. 9B). This hypothesis was guided by prior work ^43^ alluding to PST’s deleterious impact on the gut mucosa, causing macrophage infiltration and inflammation. To test the nature of contribution of PST and CST on the gut paracellular barrier function, we supplemented the CgA-KO mice with either CST alone (2 µg/g body weight for 15 days), or PST alone (2 µg/g body weight for 15 days), or both PST and CST co-administered at equimolar ratio for 15 days. CST alone did not change leakiness of FITC-dextran in CgA-KO mice (Fig. 9C).

By contrast, PST alone increased the leakiness in CgA-KO mice and this leakiness was markedly reduced when the CgA-KO mice received both PST and CST at equimolar concentration (Fig. 9C). These findings indicate that PST increases the paracellular gut permeability, and that CST antagonizes this action. Together, these data lead us to propose a working model (Fig. 9D) in which the intestinal epithelial paracellular barrier is finetuned by the EECs via a major gut hormone, CgA. Our data show that CgA is proteolytically processed into two peptides, CST and PST, which tighten and loosen the gut barrier, respectively.

## Discussion

The major finding in this work is the discovery of a molecular mechanism that involves a major EEC-derived gut hormone that has a balanced action on the paracellular gut barrier; two hormone derivatives of CgA, i.e., CST and PST, which either tighten or loosen the barrier, respectively. Because CgA-KO mice had unimpaired leakage of FITC-dextran, which was increased by exogenous administration of PST but not reduced by exogenous CST alone, findings also indicate that the barrier-homeostatic mechanism(s) is contained within and converge at a molecular level on the prohormone CgA and its processing into counteracting peptide hormones. These findings are important because CST is a major peptide hormone in EEC which is proteolytically produced from CgA by proprotein convertase 1 (PC1) ^55^, cathepsin L ^56^, plasmin ^40^ and kallikrein ^57^. Its abundance in the gut suggests that the permeability of the gut barrier is regulated by CST.

CgA is unique in having peptide domains that antagonize several functions including inflammation (CST suppresses while PST promotes), obesity (CST inhibits while PST stimulates) and sensitivity to insulin (CST increases while PST diminishes) ^20,27,58^. Since CgA-KO mice lack both CST and PST, and we observed increased permeability in CST-KO mice and decreased permeability in CgA-KO mice, we reasoned that these opposing phenotypes are probably caused by the lack of PST in the CgA-KO mice. Indeed, the increased permeability in PST-treated CgA-KO mice and its reversal by co-treatments of PST and CST lead us to conclude that CST promotes the epithelial paracellular barrier function by counteracting PST. Nevertheless, the baseline gut permeability in CST-KO mice was higher than in CgA-KO mice supplemented with PST. This might suggest that there are other cleavage fragments of CgA that regulate the intestinal permeability or that the effects cannot be completely reversed.

While PST exerts deleterious impact on the gut mucosa, causing macrophage infiltration and inflammation ^59^, we found that lack of CST caused an increased expression of proinflammatory genes and proteins and treatment of CST-KO mice with CST reversed expression of those genes and proteins. These findings are in line the anti-inflammatory roles of CST ^27,29^ and with the low-grade systemic inflammation present in the CST-KO mice ^27,29^. Our findings are also in congruence with the existing literature which showed increased TJ permeability in presence of increased proinflammatory cytokines such as TNF-α, IL-1β, and IFN-γ ^60^. A reduced gut barrier function might facilitate the leakage of microbial components such as lipopolysaccharide (LPS) and peptidoglycan (PGN) from the lumen of the gut into the tissues, where they can bind to pattern recognition receptors and activate the immune system. In turn, the increased inflammation can result in further leakage of the gut. For instance, myosin light chain kinase (MLCK) plays critical roles in cytokine-mediated TJ regulation by disrupting the interaction between TJ proteins and the actin-myosin cytoskeleton ^61^ leading to damage of the TJ scaffold, which is crucial for the maintenance of barrier integrity ^61^. Inflammatory cytokine-induced intestinal barrier loss is also believed to be due to endocytosis of TJ proteins, which also requires MLCK transcription and activity at the TJ ^6^, although we did not observe decreased protein levels of TJ proteins in the colon of CST-KO mice. In addition to the above mechanisms, TNF-α suppresses TJ barrier function by decreasing expression of TJP1 protein, increasing expression of Claudin-2, and activating the NF-κB pathway ^62^. IL-6 also increases expression of Claudin-2 ^63^. In line with this, we observed elevated levels of Claudin-2 in the colon of CST-KO mice.

In addition to these indirect effects on TJ protein expression and structure via the immune system, it might also be possible that CgA and/or its cleavage products exert direct effects on the epithelial cells and thereby more directly regulate the TJs. To investigate this, it will be interesting to perform *in vitro* experiments, e.g., epithelial monolayers in perfusion chambers co-cultured with or without immune cells ^64^.

An interesting question is how CST and PST affect the AJs and desmosomes. Not only TJ components, but also desmosome and AJ components are altered in CST-KO mice and these changes are reversible by administration of exogenous CST. It might be possible that the AJs and/or desmosomes are part of a feedback mechanism to compensate the altered TJ structure. Previously, by using Dsg2-KO mice, desmosomes have been shown to regulate the intestinal barrier ^45^. Moreover, Dsg2 modulates the production of glial cell-derived neurotrophic factor (GDNF) in inflammatory bowel disease patients ^65^. Therefore, a possible scenario is that CgA-derived peptides regulate the release of GDNF from enteric glial cells in the gut wall, which in turn affects the epithelial junctions.

The present study unequivocally demonstrates that CST is a key regulator of paracellular gut permeability since the absence of CST (CST-KO) is accompanied by dysbiosis, increased gut permeability for FITC-dextran, increased infiltration of macrophages and CD4^+^ T cells, and higher local inflammatory gene and protein expression with the substitution of CST-KO mice with CST reversing many of the above abnormalities. Do gut hormones simulate the actions of CST? While Cholecystokinin (CCK) decreases mucosal production of proinflammatory cytokines induced by LPS and increases the expression of seal forming TJ proteins (OCLN, Claudins and JAM-A) ^66^, Ghrelin ameliorates sepsis-induced gut barrier dysfunction by activation of vagus nerve via central ghrelin receptors ^53^ as well as by decreasing production of TNF-α and IL-6 ^67^. Glucagon-like peptide 1 (GLP-1) has also been shown to decrease inflammation of enterocytes ^68^. Vasoactive intestinal peptide, a neuropeptide found in lymphocytes ^69^, mast cells ^70^, and enteric neurons of the gastrointestinal tract regulates both epithelial homeostasis ^71^ and intestinal permeability ^72^. The multitude of effects of CST on gut physiology and their recapitulation by these well-investigated gut hormones establishes CST as an important gastrointestinal peptide hormone.

CST-KO mice (loss of barrier integrity) are hypertensive and insulin-resistant on normal chow diet ^27,38^ and CgA-KO mice (intact barrier integrity) have a normal blood pressure and are supersensitive to insulin in both normal chow and on high fat diet ^18,20^. Could metabolic endotoxemia explain the above phenotypes? Besides gastrointestinal peptide hormones and neuropeptides, TJ homeostasis is altered by proinflammatory cytokines, pathogenic bacteria, and pathological conditions like insulin resistance and obesity. Firmicutes (e.g., *Lactobacillus, Clostridium*, and *Enterococcus*) and Bacteroidetes (e.g., *Bacteroides*) constitute the major bacterial phyla in the gut ^73^. Toxigenic bacteria cause diarrhea *via* increased ion permeability of the pore pathway ^51^, but gut microbes might also affect the leak pathway. Enteric pathogenic bacteria such as *Escherichia coli* and *Salmonella typhimurium* alter the intestinal epithelial TJ barrier and cause intestinal inflammation ^74^. LPS alters expression and localization of TJ proteins such as TJP1 and OCLN ^75^. Therefore, it would be interesting to study whether the systemic LPS levels support the observed dysbiosis-related loss of gut barrier in the CST-KO mice. Since high fat diet-induced insulin-resistant mice and patients with obesity or diabetes display elevated levels of Firmicutes and Proteobacteria compared to the beneficial phylum Bacteroidetes ^76^, it is conceivable that dysbiosis (i.e., increased Firmicutes and decreased Bacteroidetes) in CST-KO mice is due to their resistance to insulin. This is further supported by the fact that *Akkermansia muciniphila*, which regulates intestinal barrier integrity, is less abundant in diabetic individuals ^77^. Furthermore, the daily administration of *A. muciniphila* is known to mitigate high fat-induced gut barrier dysfunction ^78^. Conversely, metabolic endotoxemia, resulting from the loss of barrier integrity, contributes to insulin resistance and obesity ^76,79^. The intact paracellular barrier function in CgA-KO mice could thus be due to their heightened sensitivity to insulin.

Human calprotectin, a 24 kDa dimer, is formed by S100A8 and S100A9 monomers and released from neutrophils and monocytes. In fact, this calprotectin complex constitutes up to 60% of the soluble proteins in the cytosol of neutrophils ^80^ and fecal calprotectin levels serve as a marker of intestinal inflammation ^81,82^. Because of the positive correlation between calprotectin concentration in gut lavage fluid and intestinal permeability ^83^, calprotectin levels are also used to assess intestinal permeability. Increased calprotectin levels have been reported to increase intestinal permeability ^49^. Therefore, increased gene and protein expressions in the colon of CST-KO mice are in line with the increased permeability in CST-KO mice.

Elevated plasma CST levels in patients suffering from Crohn’s disease both in disease remission and flare-up as compared to elevated precursor CgA in Crohn’s disease flare-ups only suggests that while CgA is generally upregulated in Crohn’s disease, it is only efficiently processed to CST during disease remission. Previous studies revealed higher plasma CgA levels during flare-ups as compared to ulcerative colitis and Crohn’s disease patients in remission ^34^. An open question remains how the levels of PST and CST are regulated in ulcerative colitis and Crohn’s disease patients. For CST our findings suggest a proteolytic switch, where CgA and CST are excessively produced in Crohn’s disease regardless of the disease state, but CgA is converted more into the anti-inflammatory CST in quiescent Crohn’s disease. The gut barrier dysfunction in CST-KO mice is reminiscent of active Crohn’s disease, because the gut permeability provided by cell-cell junctions is generally diminished in IBD patients ^84^. Based on our findings, we hypothesize that the higher levels of CST during Crohn’s disease remision promote anti-inflammatory responses and aid restoration of epithelial cellular junctions.

## Material and Methods

### Mice

Male wild type (WT), CST-knockout (KO) and CgA-KO mice (3 weeks old) on C57BL/6 background were kept in a 12 hr dark/light cycle on normal chow diet (NCD: 13.5% calorie from fat; LabDiet 5001, Lab Supply, Fort Worth, TX). Experiments were conducted in a blinded fashion whenever possible. For terminal procedure, mice were deeply anesthetized (assessed by pinching toe response) with isoflurane followed by harvesting of tissues. Euthanasia was achieved through exsanguination. For rescue experiments with exogenous CST and PST, mice were injected intraperitoneally with CST (2 µg/g body weight) and/or PST (2 µg/g body weight) at 9:00 AM for 2-4 weeks before collecting feces or harvesting tissues. All mouse studies were approved by the UCSD and Veteran Affairs San Diego Institutional Animal Care and Use Committees and conform to relevant National Institutes of Health guidelines.

### Gut permeability assay

Fluorescein isothiocyanate conjugated dextran (FITC-dextran-FD4, 4 kDa; MilliporeSigma, St. Louis, MO) was administered (44 mg/100 g body weight) to 8 hr fasting mice by oral gavage followed by collection of blood after 4 hr of administration from the heart under deep isloflurane anesthesia ^85^. Plasma concentration of FITC was determined by comparison of fluorescence signals with a FITC-dextran standard curve.

### Measurement of tissue cytokines, CST, CgA and calprotectin

A portion of the colon was homogenized in PBS and cytokines were measured in 20 µl of the homogenate using U-PLEX mouse cytokine assay kit (Meso Scale Diagnostics, Rockville, MD) via the manufacturer’s protocol. Mouse EIA kits from CUSABIO (CUSABIO Technology LLC, Houston, TX) were used to determine CgA (CSB-EL005344MO), CST (CSB-E17357m), and calprotectin (CSB-EQ013485MO).

### Determination of CgA and CST levels in human plasma

EDTA-plasma was collected from patients with Crohn’s disease (N=89) and ulcerative colitis (N=101) (Supplemental table S1) and healthy controls (N=50). Intestinal inflammation was assessed by colonoscopy. A flare-up was defined as histological and endoscopic disease activity, whereas remission entailed absence of endoscopic inflammation and histological disease activity. EDTA-plasma from healthy controls (Supplementary table S2) was obtained from the Mini Donor Service at the University Medical Center Utrecht. Samples were thawed and diluted 50-times with extraction buffer (PBS with 0.5% Tween and 0.05% sodium azide), thoroughly vortexed followed by incubation at 4°C for 20 min on a rolling device. Afterwards samples were centrifuged at 760 ×g for 20 min at 4°C and the supernatant was collected for analysis. Samples were collected in compliance with the Declaration of Helsinki. EDTA plasma was used to quantify CgA and CST using ELISA (CUSABIO CSB-E17355h, CSB-E09153h). Informed consent was obtained, and the study was conducted in accordance with the Institutional Board Review of the University Medical Center Utrecht (approval number NL35053.041.11 &11-050).

### Colon histological stains and immunohistochemistry

Mouse colon was isolated, washed in PBS and dehydrated in ethanol series, xylene and paraffin. Afterwards, the tissue was embedded in paraffin, formalin fixed, and paraffin embedded (FFPE) and sections of 5 µM were cut. Hematoxylin and eosin (HE) and Masson’s trichrome (MT) staining were performed using standard techniques. For the immunohistochemistry, the tissue was heated in citrate buffer (pH 6.0, 1.0 nM citric acid) and endogenous peroxidase activity was quenched with 3% H_2_O_2_ in methanol. Standard indirect immunoperoxidase procedures were used for detection (Vectastain and SK-4800, Vector Laboratories) with the antibody Rabbit-αKi-67 (1:100, Novus Biologicals), Rat-αF4/80 (1:500, Caltag laboratories and slides were counterstained with Mayer’s haematoxylin. Followed by mounting using Quick D mounting medium. For fluorescent staining, slides were incubated with Rabbit-αCASPIII antibody (1:100, ab2302, Abcam) and nuclear counterstain with DAPI using Fluoromount-G (0100-01, Southern Biotech). Slides were imaged using the PerkinElmer Vectra (Vectra 3.0.3; PerkinElmer, MA) at 20x magnification or imaged using the Leica SP8 with a water dipping objective.

### Transmission Electron Microscopy (TEM) and morphometric analysis

To displace blood and wash tissues before fixation, mice were cannulated through the apex of the heart and perfused with a Ca^2+^ and Mg^2+^ free buffer composed of DPBS and 9.46 mM KCl as described ^73^. Mouse and human colon fixation, embedding, sectioning and staining were done as described ^73^. Grids were viewed using a JEOL JEM1400-plus TEM (JEOL, Peabody, MA) and photographed using a Gatan OneView digital camera with 4×4k resolution (Gatan, Pleasanton, CA). Morphometric analyses of the dense core vesicles were determined as described ^86^. TJ length and diameter were determined by measuring the lengths and perpendicular widths using the line tool in NIH ImageJ software.

### Tight junction protein analysis via immunoblotting

Mouse colon pieces were homogenized in a buffer containing phosphatase and protease inhibitors, as previously described ^19^. Colon lysates were subjected to SDS-PAGE and immunoblotted. Followed by staining with Rabbit-αOccludin (1:1000, ab216327, Abcam, Cambridge, MA, USA), Rabbit-αClaudin 1 (1:2000, ab180158, Abcam), Rabbit-αClaudin 2 (2µg/ml, 51-6100, Thermo Fisher Scientific, Waltham, MA, USA), Mouse-αE-cadherin (1:500, 610182, BD Biosciences, San Jose, CA, USA), Rabbit-αcatenin (1:1000, C2081, Sigma-Aldrich, St. Louis, MO, USA) or Rat-αZO-1 (1:500, R26.4C, DSHB, Iowa City, Iowa, USA).

### Flow cytometry analysis

Isolated colon cells from digested mouse colon were stained with fluorescence-tagged antibodies to detect macrophages (CD45^+^CD11b^+^Emr1^+^) and helper T cells (CD45^+^CD4^+^). Data were analyzed using FlowJo.

### Microbiome analysis

DNA was extracted from cecum of WT, CST-KO and CgA-KO (normal chow diet (NCD), littermates) using the Zymobiomics miniprep kit. Isolated DNA was quantified by Qubit (Thermo Fisher Scientific). DNA libraries were prepared using the Illumina Nextera XT library preparation kit. Library quantity and quality was assessed with Qubit and Tapestation (Agilent Technologies). Libraries were sequenced on Illumina HiSeq platform 2×150bp. Unassembled sequencing reads were directly analyzed by CosmosID bioinformatics platform (CosmosID) ^87^ for multi-kingdom microbiome analysis and profiling of antibiotic resistance and virulence genes and quantification of organism’s relative abundance. This system utilizes curated genome databases and a high-performance data-mining algorithm that rapidly disambiguates hundreds of millions of metagenomic sequence reads into the discrete microorganisms engendering the particular sequences. Data mining, interpretation and comparison of taxonomic information from the metagenomic datasets were performed using Calyspo Bioinformatics software V.8.84.

### Real Time PCR

Total RNA from colon tissue was isolated using RNeasy Mini Kit and reverse-transcribed using qScript cDNA synthesis kit. cDNA samples were amplified using PERFECTA SYBR FASTMIX L-ROX 1250 and analyzed on an Applied Biosystems 7500 Fast Real-Time PCR system. Primer sequences are in Supplemental table S3. All PCRs were normalized to *Rplp0*, and relative expression levels were determined by the ΔΔ*C*_*t*_ method as described ^24^.

### Statistics

Statistics were performed with PRISM 8 (version 8.4.3; San Diego, CA). Data were analyzed using either unpaired two-tailed Student’s *t*-test for comparison of two groups or one-way or two-way analysis of variance for comparison of more than two groups followed by Tukey’s *post hoc* test if appropriate. All data are presented as mean ± SEM and significance were assumed when p<0.05.

## Supporting information

Supplementary tables MS CST-KO gut permeability

## Declarations

## Acknowledgements

We thank Zbigniew Mikulski (La Jolla Institute for Immunology, CA, USA) and Kiek Verrijp (Radboud University Medical Center, Netherland) for technical support with the histology and immunohistochemistry. This work was supported by grants from the Veterans Affairs (I01 BX003934 to SKM), the Human Frontier Science Program (HFSP; RGY0080/2018 to G.v.d.B), the Netherlands Organization for Scientific Research (NWO-ALW VIDI 864.14.001 to G.v.d.B), and European Research Council (grant agreement No. 862137 to G.v.d.B). G.C. is supported by grants from the Swedish Research Council and the Swedish Society for Medical Research. S.E.A. is funded by is supported by a Rosalind Franklin Fellowship, co-funded by the European Union and University of Groningen, The Netherlands. E.M.M is supported by a short-term EMBO fellowship (EMBO7887).

## Conflicts of interest

The authors declare no conflict of interest.

## Availability of data and material

Primary data available from the corresponding authors on reasonable request.

## Notes

### Competing Interest Statement

The authors have declared no competing interest.

### Summary of Updates

We have revised the original figures with new data and also included new data in new figures. The revised figures on the expression of genes include WT+CST-treated data (Fig. 2F-H, 4G and 5C). The new figures include the following: Western blots of TJ and AJ proteins (Fig. 3A and B); ultrastructural analyses of human colonic mucosa (Fig. 3H-L); histology showing immune cells (Fig. 4A-C); immunohistochemistry (Emr+ staining; Fig. 4D); Masson trichrome staining showing fibrosis (Fig. 5D), transmission electron micrographs showing TJ, AJ and desmosomes in healthy volunteers and IDB patients (Fig. 2H-L) as well as collagen fibers (Fig. 5E), and immunohistochemistry of caspase 3 showing apoptosis (Fig. 6A).

