## Supplementary tables MS CST-KO gut permeability for "Chromogranin A regulates gut permeability *via* the antagonistic actions of its proteolytic peptides"

[1]Department of Tumor Immunology, Radboud Institute for Molecular Life Sciences, Radboud University Medical Center, Nijmegen, the Netherlands; [2]VA San Diego Healthcare System, 3350 La Jolla Village Drive, San Diego, CA, USA; [3]Center for Translational Immunology, Utrecht University Medical Center, Utrecht, the Netherlands; [4]Department of Gastroenterology and Hepatology, Utrecht University Medical Center, Utrecht, the Netherlands; [5]Science for Life Laboratory and [6]Department of Medical Cell biology, Uppsala University, Uppsala, Sweden; [7]Departments of Cellular and Molecular Medicine, [9]Pathology, [10]Medicine, University of California San Diego, La Jolla, CA, USA; [11]Department of Molecular Immunology and Microbiology, Groningen Biomolecular Sciences and Biotechnology Institute, University of Groningen, Groningen, the Netherlands.

*shared co-first authorship

**Short title:** Catestatin regulates gut permeability

**Supplemental Table S1. Patient characteristics.**[1]

| | **Healthy controls** (n = 50) | **CD remission** (n = 49) | | **CD flare** (n = 40) | **UC remission** (n = 51) | | **UC flare** (n = 50) |
|---|---|---|---|---|---|---|---|
| **Female** (n, %) | 41 (82%) | 32 (65%) | | 25 (51%) | 24 (47%) | | 23 (46%) |
| **Age** (average, range) | 43 (21-68) | 49 (22-71) | | 46 (20-72) | 51 (20-76) | | 49 (24-75) |
| **Disease duration** (average, range) | - | 21 (8-51) | | 18 (7-48) | 19 (6-46) | | 19 (3-51) |
| **Montreal** (n, %) | - | *L1* | 2 (4%) | 0 (0%) | *E1* | 16%) | 2 (4%) |
| | | *L2* | 21 (43%) | 18 (37%) | *E2* | 20 (39%) | 14 (28%) |
| | | *L3* | 24 (49%) | 28 (57%) | *E3* | 22 (41%) | 34 (68%) |
| | | *L2 + L4* | 0 (0%) | 2 (4%) | | | |
| | | *L3 + L4* | 2 (4%) | 1(2%) | | | |
| **IBD medication** *(%)* *None* | 50 (100%) | 9 (18%) | | 14 (29%) | 7 (14%) | | 1 (2%) |
| *Mesalazine preparations* | | 9 (18%) | | 5 (10%) | 36 (71%) | | 39 (78%) |
| *Corticosteroids* | | 1 (2%) | | 6 (12%) | 0 (0%) | | 7 (14%) |
| *Immunosuppressives* | | 26 (53%) | | 14 (29%) | 20 (39%) | | 15 (30%) |
| *Biologicals* | | 12 (24%) | | 16 (33%) | 4 (8%) | | 3 (6%) |
| **TNFα plasma levels** (nM) (average, range) | 5.9 (0-21.9) | 2.6 (0-15.8) | | 5.2 (0-26.1) | 3.4 (0-33.8) | | 5.5 (0-45.6) |

[1]Characteristics of the patients at time of inclusion. As an indication for the extent of disease, the location has been scored according to the Montreal classification. For CD; L1 is ileal disease, L2 is colonic disease, L3 is ileocolonic disease, and L4 is isolated upper gastro-intestinal tract. For UC; E1 is distal to the rectosigmoid junction (proctitis), E2 is distal to the splenic flexure (left-sided) and E3 is proximal to the splenic flexure (extensive). Medication entails therapeutic categories in use for IBD; other therapies are not included. Some patients use more than one drug which all have been scored separately. Biologicals comprised anti-TNF-α compounds except for one patient in the CD flare group. CD: Crohn's disease, IBD: inflammatory bowel disease, UC: ulcerative colitis, TNF-α: tumor necrosis factor alpha. No correlation was observed between Chromogranin A and TNF-α levels (Pearson's correlation: $r = 0.12$, $p = 0.08$). Also no correlation was observed between Chromogranin A and TNF-α levels when plotted per disease group (healthy control, UC and CD remission and flare-up; Table S2).

**Supplemental Table S2. Correlation matrix patient characteristics.**[1]

| | **CgA** (nM) | **CST** (nM) | **TNFα** (nM) |
|---|---|---|---|
| **CgA** (nM) | 1.000 | 0.221 | 0.127 |
| **CST** (nM) | 0.221 | 1.000 | -0.002 |
| **TNFα** (nM) | 0.127 | -0.002 | 1.000 |
| **Disease status** | 0.196 | 0.298 | 0.038 |
| **Sex** | 0.020 | -0.147 | -0.111 |
| **Age** | -0.006 | -0.008 | -0.066 |
| **Years since diagnosis** | -0.143 | 0.006 | 0.027 |
| **Montreal classification** | 0.030 | -0.034 | 0.057 |
| **Clinical symptoms** | 0.147 | -0.052 | 0.250 |
| **Medication use** | -0.079 | -0.118 | 0.093 |
| **Mesalazine preparations** | -0.009 | -0.136 | 0.003 |
| **Corticosteroids** | 0.079 | 0.077 | 0.102 |
| **Immunosuppressives** | -0.062 | 0.016 | -0.028 |
| **Biologicals** | 0.059 | -0.086 | 0.064 |
| **anti-TNFα therapy** | 0.069 | -0.051 | 0.030 |

[1]Correlation matrix of all patient characteristics (removed endoscopy and pathology due to many absent values, and healthy control subset as most factors were not determined). CgA and CST correlated strongest with disease status (CD and UC remission and flare-up). All characteristics were categorical (yes/no or female/male) expect for age, years since diagnosis and CgA, CST and TNF-α levels. A log transformation was performed when a characteristic was not normally distributed.

**Supplemental Table S3. Primer sequences for RT-qPCR experiments.**

| Gene | Gene ID | Forward primer (5'-3') | Reverse primer (3'-5') |
|---|---|---|---|
| *Arg1* | 11846 | CTCCAAGCCAAAGTCCTTAGAG | AGGAGCTGTCATTAGGGACATC |
| *Ccl2* | 20296 | TTAAAAACCTGGATCGGAACCAA | GCATTAGCTTCAGATTTACGGGT |
| *Cdh1* | 12550 | AGGAAATGCACCCCTCCAAT | AATCGGCCAGCATTTTCTG |
| *Chga* | 12652 | AGCCAGACTACAGACCCACT | TGACTTCCAGGACGCACTTC |
| *Cldn1* | 12737 | GGGGACAACATCGTGACCG | AGGAGTCGAAGACTTTGCACT |
| *Cldn2* | 12738 | CAACTGGTGGGCTACATCCTA | CCCTTGGAAAAGCCAACCG |
| *Cldn6* | 54419 | ATGGCCTCTACTGGTCTGCAA | GCCAACAGTGAGTCATACACCTT |
| *Clec7a* | 56644 | GACTTCAGCACTCAAGACATCC | TTGTGTCGCCAAAATGCTAGG |
| *Clec10a* | 17312 | TGAGAAAGGCTTTAAGAACTGGG | GACCACCTGTAGTGATGTGGG |
| *Ctnna1* | 12385 | ACTTTGATGTCAGAAGCAGGACC | CACCTGTTCTGCAATCTTTGCTTT |
| *Ctnnb1* | 12387 | AAGGAAGCTTCCAGACATGC | AGCTTGCTCTCTTGATTGCC |
| *Ctsl* | 13039 | ATCAAACCTTTAGTGCAGAGTGG | CTGTATTCCCCGTTGTGTAGC |
| *Cxcl1* | 14825 | GCTTGAAGGTGTTGCCCTCAG | AAGCCTCGCGACCATTCTTG |
| *Dsg2* | 13511 | CGTGGTTGAAGGCATTCATTTC | TAGCTGCTTGACCAGTGTCTT |
| *Emr1* | 13733 | TTGTACGTGCAACTCAGGACT | GATCCCAGAGTGTTGATGCAA |
| *F11r* | 16456 | TCTCTTCACGTCTATGATCCTGG | TTTGATGGACTCGTTCTCGGG |
| *Ifng* | 15978 | ATGAACGCTACACACTGCATC | CCATCCTTTTGCCAGTTCCTC |
| *Il10* | 16153 | GCTCTTACTGACTGGCATGAG | CGCAGCTCTAGGAGCATGTG |
| *Il12b* | 16160 | CCCTGACATTCTGCGTTCA | AGGTCTTGTCCGTGAAGACTCTA |
| *Il4* | 16189 | GGTCTCAACCCCCAGCTAGT | GCCGATGATCTCTCTCAAGTGAT |
| *Itgam* | 16409 | ATGGACGCTGATGGCAATACC | TCCCCATTCACGTCTCCCA |
| *Itgax* | 16411 | CTGGATAGCCTTTCTTCTGCTG | GCACACTGTGTCCGAACTCA |
| *Marveld2* | 218518 | GGGTCGCAAGGCACCTTTAAT | ACCTTCTCAGAATACGTCCGC |
| *Mrc1* | 17533 | CTCTGTTCAGCTATTGGACGC | CTCTGTTCAGCTATTGGACGC |
| *Nos2* | 18126 | GTTCTCAGCCCAACAATACAAGA | GTGGACGGGTCGATGTCAC |
| *Ocln* | 18260 | TTGAAAGTCCACCTCCTTACAGA | CCGGATAAAAAGAGTACGCTGG |
| *Plg* | 18815 | TGCAGTGGAGAAAAGTATGAGGG | AGGGATGTATCCATGAGCATGT |
| *Rplp0* | 11837 | AGATTCGGGATATGCTGTTGGC | TCGGGTCCTAGACCAGTGTTC |
| *S100a8* | 20201 | AAATCACCATGCCCTCTACAAG | CCCACTTTTATCACCATCGCAA |
| *S100a9* | 20202 | ATACTCTAGGAAGGAAGGACACC | TCCATGATGTCATTTATGAGGGC |
| *Tgfb1* | 21803 | CTCCCGTGGCTTCTAGTGC | GCCTTAGTTTGGACAGGATCTG |
| *Tjp1* | 21872 | GCCGCTAAGAGCACAGCAA | TCCCCACTCTGAAAATGAGGA |
| *Tjp2* | 21873 | ATGGGAGCAGTACACCGTGA | TGACCACCCTGTCATTTTCTTG |
| *Tjp3* | 27375 | GGATGAGATCTTGCAGGTGAATGG | TCCTGTTTGCTCTGTGTTACCAGCTC |
| *Tnf* | 21926 | CCCTCACACTCAGATCATCTTCT | GCTACGACGTGGGCTACAG |
